# QSMxT: Robust Masking and Artefact Reduction for Quantitative Susceptibility Mapping

**DOI:** 10.1101/2021.05.05.442850

**Authors:** Ashley Wilton Stewart, Simon Daniel Robinson, Kieran O’Brien, Jin Jin, Georg Widhalm, Gilbert Hangel, Angela Walls, Jonathan Goodwin, Korbinian Eckstein, Monique Tourell, Catherine Morgan, Aswin Narayanan, Markus Barth, Steffen Bollmann

## Abstract

**Purpose:** Quantitative Susceptibility Mapping (QSM) is a post-processing technique applied to gradient-echo phase data. QSM algorithms require a signal mask to delineate regions with reliable phase signal for subsequent susceptibility estimation. Existing masking techniques used in QSM have limitations that introduce artefacts, exclude anatomical detail, and rely on parameter tuning and anatomical priors that narrow their application. Here, a robust masking and reconstruction procedure is presented to overcome these limitations and enable automated QSM processing for a wide range of use-cases implemented in an open-source software framework: *QSMxT*.

**Methods:** A robust masking technique that automatically separates reliable from less reliable phase regions was developed and combined with a two-pass reconstruction procedure that operates on the separated sources before combination, extracting more information while reducing the influence of artefacts.

**Results:** Compared with standard masking and reconstruction procedures, the two-pass inversion reduces streaking artefacts caused by unreliable phase and high dynamic ranges of susceptibility sources. QSMxT is robust across a range of datasets at 3 T in healthy volunteers and phantoms, at 7 T in tumour patients, and in the QSM challenge 2.0 simulated brain dataset, with significant artefact and error reductions, greater anatomical detail, and minimal parameter tuning.

**Conclusion:** QSMxT generates masks for QSM that separate reliable from less reliable phase regions, enables a more accurate QSM reconstruction that mitigates artefacts, operates without anatomical priors, and requires minimal parameter tuning. QSMxT makes QSM processing more accessible, reliable and reproducible.

## 1 Introduction

Quantitative Susceptibility Mapping (QSM) is an MRI method that aims to calculate the magnetisation induced in tissues when placed in a magnetic field. Recovering a magnetic susceptibility distribution requires a series of post-processing techniques applied to gradient-echo (GRE) phase data [1]. Compared with related techniques currently used in clinical brain imaging such as Susceptibility Weighted Imaging (SWI) [2], QSM provides improved detection and unambiguous differentiation of iron and calcium deposits, enhanced characterisation of pathological iron and myelin variations, and unique brain morphology detail within deep brain nuclei [3, 4]. QSM applications include brain imaging to study Huntington’s disease [5], multiple sclerosis [6–9], Alzheimer’s disease [10, 11] and Parkinson’s disease [12, 13], as well as applications outside the brain such as quantifying liver iron [14, 15], identifying calcifications in the prostate [16] and imaging cartilage in joints [17].

QSM reconstruction classically involves several independent phase processing steps, including phase combination from multiple coils, phase unwrapping, background field removal, and dipole inversion [1]. Phase combination determines the inhomogeneous coil-dependent phase offsets, which are necessary to produce a corrected phase signal that reflects the *B*_0_ field [18]. Phase unwrapping removes the aliasing that occurs in measured phase values as they wrap into the range (™π, π] [18]. Background field removal estimates and subtracts contributions to the phase that originate from susceptibility sources outside the region of interest (ROI) [19]. Finally, the dipole inversion step recovers the magnetic susceptibility by applying an inverse physical model to the sample-induced phase shifts [1, 3]. Some QSM processing techniques combine several of these steps into a single optimisation problem, improving computational efficiency and reducing error propagation [20], or in more recent work, learn the inverse problem using convolutional neural networks [21].

In practice, QSM reconstruction involves several other processing steps. First, scanner-produced images (such as DICOMs) are converted to processing formats (such as NIfTI) [22, 23]. A mask is generated or drawn to delineate the ROIs containing reliable phase values prior to susceptibility calculation [3]. Multi-echo data are combined across echoes [24] through an additional phase combination step [25], or by averaging the QSM result across echoes. Finally, studies involving multiple participants require the construction of a common space to facilitate group-level analyses, as well as segmentation of the susceptibility maps to extract quantitative values in anatomical regions of interest. These segmentations can be manually drawn or generated automatically, usually via a T1-weighted scan registered to the QSM space.

For most publicly available QSM processing pipelines, these pre- and post-processing tasks are out-of-scope, with a strong focus only on one or more of the phase unwrapping, background field removal and dipole inversion steps [26–28], and thus manual intervention for the additional steps is generally required. Using such tools across large-scale datasets without automation is laborious and infeasible for clinical QSM users, and a hurdle to researchers less familiar with QSM. A fully automated solution requires the configuration of a wide range of software tools, making it challenging to setup. However, recent developments in the field, such as the SEPIA toolbox [29], attempt to provide extensible frameworks for QSM processing, enabling a wide range of algorithms to be supported by a single and easy-to-use tool, though several steps are unaccounted for, manual intervention is required, and processing does not scale well for large datasets.

One critical step that is often neglected in the QSM literature is signal masking. Signal masks in QSM are necessary to identify the regions containing reliable phase signal for the background field removal and dipole inversion steps. Including unreliable voxels introduces errors in the background field estimation and erroneously strong susceptibility values as well as streaking artefacts throughout the final susceptibility map. Most QSM literature cites the Brain Extraction Tool (BET) from the FMRIB Software Library (FSL) for mask generation [30], as the QSM algorithms themselves do not generate a mask or specify their requirements. BET segments brain from non-brain in the signal magnitude by applying locally adaptive forces to a deformable model, evolving it to fit the brain surface. Using a brain mask for QSM has several limitations that impact reconstruction quality and utility. Firstly, areas with high SNR such as the medulla and lower brainstem are specifically excluded, which is a problem for studies investigating regions such as the neck [31]. Secondly, the assumption of normal brain anatomy for masking is not always warranted and impacts the application of QSM in patients with brain tumours or traumatic brain injuries [32, 33]. Another tenuous assumption is that all brain voxels contain sufficiently reliable phase for QSM despite rapid signal decay with increasing echo time (TE), especially near the brain boundary and strong susceptibility sources such as vessels and lesions. Reconstructing QSM in such areas and across high dynamic ranges of susceptibility values is challenging, with most reconstruction algorithms introducing pervasive streaking artefacts without sufficient regularisation. However, strong regularisation causes loss of detail throughout the susceptibility map. Despite this, and even in multi-echo acquisitions, a single brain mask is often used, ignoring the behaviour of the phase signal and potentially introducing artefacts, especially at later TEs.

Recent QSM algorithms such as STAR-QSM [34], QSMART [35], and superposed dipole inversion (SDI) [36] address the high dynamic range problem by resolving weak and strong susceptibility sources in separate inversion steps so that artefacts are avoided or specifically excluded after a combination step. STAR-QSM reconstructs strong sources first using a high regularisation factor before simulating the phase shifts caused by these sources using the dipole forward model. These phase shifts are then subtracted from the original phase so that subtle sources can be reconstructed in the second pass with a lower regularisation factor before combination. QSMART, on the other hand, uses a different mask for each inversion, and both can be performed in parallel. The first reconstruction uses a full brain mask, while the second uses a brain mask with vasculature removed. Vasculature is identified by applying a Frangi vessel enhancement filter on the echo-averaged magnitude data. The remaining method, SDI, performs a single QSM reconstruction using a brain mask, before generating a second mask by thresholding the QSM result to remove strong, positive susceptibility sources. The second mask is used for a second QSM reconstruction, with both results combined by superposition. The currently existing QSM techniques have limitations, as they use masks generated using anatomical priors for brain imaging, and require application-specific regularisation parameter tuning. These limitations narrow the application potential for each method and reduce their robustness across diverse datasets.

In this paper, a robust masking technique and two-pass reconstruction procedure are presented as part of the open-source QSM framework, *QSMxT* [37]. Two-pass QSM aims to mitigate streaking artefacts throughout susceptibility maps using the signal intensity or phase coherence to separate reliable from less reliable sources [34, 35, 38]. Since the technique is based on thresholding, it does not require anatomical priors, is easily automated, and is robust to pathology. These features enable users to explore many applications otherwise requiring bespoke or cumbersome manual techniques impractical for large datasets. The automation potential of two-pass QSM is demonstrated by its integration within a fully automated QSM framework, QSMxT. QSMxT automates all steps from raw data preparation to QSM reconstruction, group space and template generation, as well as region of interest analyses across large datasets, along with all required dependencies. Support for parallel processing on multi-core or cluster computing systems ensures QSMxT is both practical and scalable, and its availability on every operating system via a modular and containerised ecosystem makes it reproducible and deployable.

## 2 Methods

### 2.1 Data and acquisition

GRE datasets from 46 healthy volunteers were acquired with 1 mm isotropic resolution, TE=5.84 /10.63/15.42/20.21/25.00 ms, TR=29.9, FA=15°, flow compensation enabled, phase encoding in the row direction, and pixel bandwidth=310.0 Hz. T1-weighted datasets were also acquired with 1 mm isotropic resolution and TE=2.96 ms. Data were acquired using a MAGNETOM Skyra 3T scanner (Siemens Healthcare GmbH, Erlangen, Germany) with a 64-channel head/neck coil on software version VE11. Phased array data were combined using the Pre-scan Normalise plus Adaptive Combine approach [39].

GRE datasets from 6 brain tumour patients were acquired with 0.3 × 0.3 × 2.9 mm resolution, TE=3.31/6.62/9.93/12.49/15.05/17.61/20.17/22.73/25.29/27.85 ms, TR=31.0 ms, FA=10°, flow compensation disabled, phase encoding in the row direction, and pixel bandwidth=523.0 Hz. Data were acquired using a MAGNETOM whole-body 7T scanner (Siemens Healthcare GmbH, Erlangen, Germany) with a 32-channel Rx head array coil (Nova Medical, Wilmington, MA, USA), on software version VB17. Phased array data were combined using ASPIRE [40].

A GRE dataset of a gel phantom containing tooth pieces and gold fiducial markers was also acquired. The phantom was 12 × 8 × 3 cm in size and made from Liquid Soft Plastic (U-Make-Em Soft Plastics, Australia) poured into a plastic container before heating to solidify. Two gold seeds were placed into the phantom, including a pure gold marker (Riverpoint Medical, Portland, Oregon, United States) measured at 1 × 3 mm, and a Gold Anchor MR+™ with 1.5% iron (Naslund Medical, Huddinge, Sweden), measured at 0.4 × 0.2 mm. The dataset was acquired with 1.1 × 1.1 × 1.2 mm resolution, TE=3.69/7.40/11.11/14.82 ms, TR=25.0 ms, FA=45.0°, flow compensation disabled, phase encoding in the row direction, and pixel bandwidth=400.0 Hz. Data were acquired using a MAGNETOM Skyra 3T scanner (Siemens Healthcare GmbH, Erlangen, Germany) with a 32-channel spine coil on software version VE11. Phased array data were combined using the Pre-scan Normalise plus Adaptive Combine approach [39].

The effect of strong susceptibility sources was further investigated using the simulated brain dataset from the QSM Challenge 2.0, which includes a simulated intracranial calcification [41, 42]. The simulation is based on a segmented MP2RAGE dataset with 0.64 mm resolution down-sampled to 1mm using k-space cropping with simulated echo-times TE=4/12/20/28 ms.

### 2.2 QSMxT Architecture

A modular reconstruction and analysis ecosystem that includes a variety of open-source tools for computing and analysing QSM data was developed. The first module consists of a continuous integration system using GitHub actions to automatically build software containers, including the container required for the QSMxT framework. The QSMxT container incorporates the FMRIB Software Library (FSL) v6.0.4 [43], FreeSurfer (v7.1.1) [44–46], Julia v1.5.3 [47], Bidscoin v2.3 [48], Python v3.8 [49], TGV-QSM v1.0.0 [20, 28], the Python packages pydicom [50], nibabel [51], seaborn [52, 53] and NiPype [54], the Julia packages ArgParse [55] and MriResearchTools [56], as well as the QSMxT NiPype workflow [37]. QSMxT runs natively on Linux operating systems and High-Performance Clusters (HPCs), only requiring Singularity [57] as a dependency. A lightweight Linux desktop is also provided, (https://github.com/NeuroDesk/vnm/) conveniently accessible via a browser interface that runs on Windows, MacOS, and Linux, only requiring Docker as a dependency.

QSMxT first converts DICOM data to NIfTI format, following Brain Imaging Data Structure (BIDS) conventions where specified [48]. Then, the NiPype QSM pipeline processes images from each subject from the BIDS directory to produce QSMs (including a combined multi-echo QSM). If T1-weighted images are available, an additional NiPype pipeline segments these and registers them to the QSM space for each subject and exports susceptibility statistics for each segmented ROI. A final NiPype pipeline generates and registers all subjects to a common group space and produces QSM and magnitude templates using minimum deformation averaging [58] (see figure 1).

**Figure 1:**
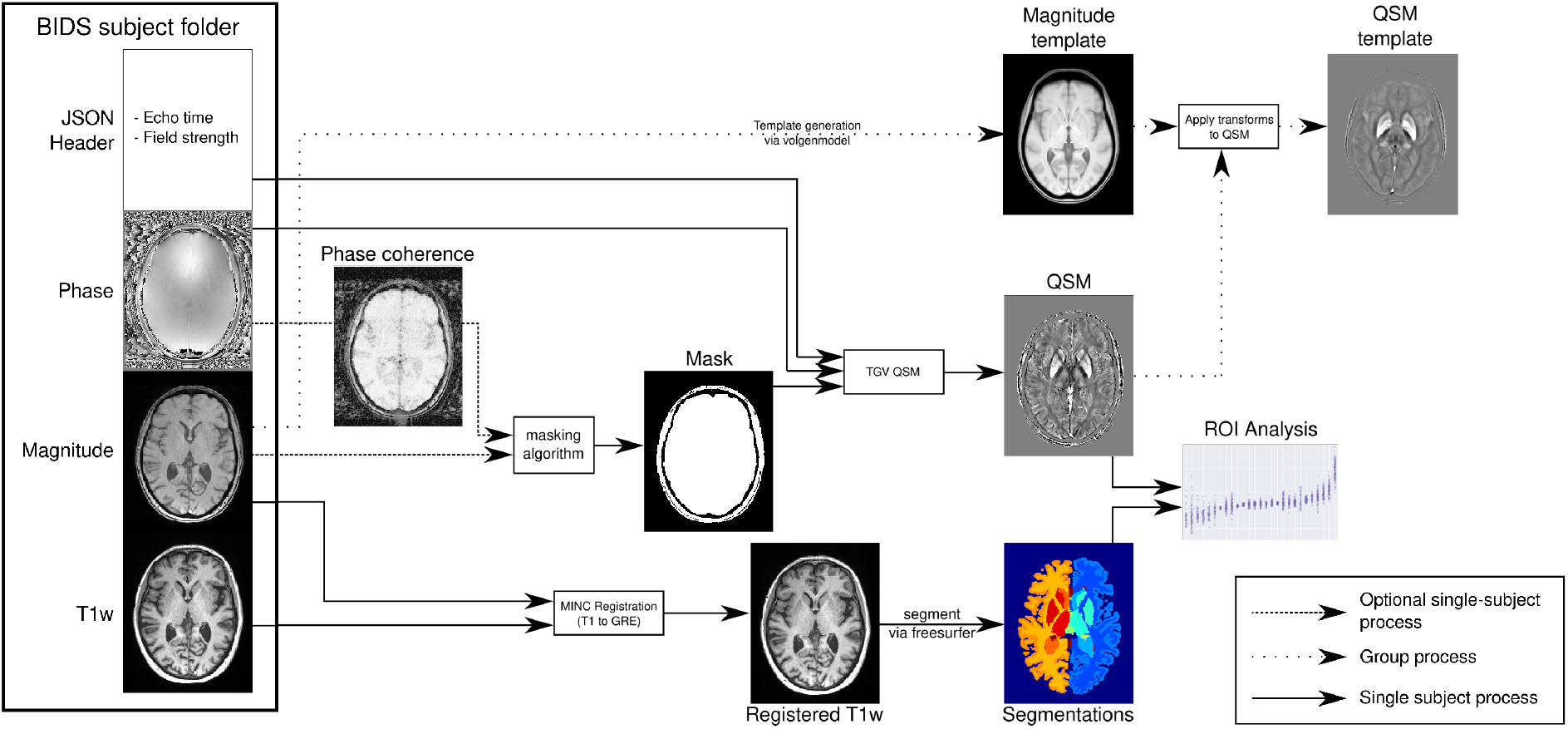
QSMxT processing pipeline and outputs after automated data conversion to the BIDS standard. Outputs for each subject include masks, T1-weighted images, segmentations registered to the QSM space, QSM images, as well as data and figures for quantitative analysis. Outputs for the whole dataset include magnitude and QSM templates.

### 2.3 QSM Reconstruction and Two-Pass QSM

The phase unwrapping, background field removal and dipole inversion steps are achieved using the *TGV-QSM* algorithm [20]. QSM algorithms incorporating TGV have been shown to be quantitatively stable across echo times compared to others [59], and achieved competitive results in the QSM Challenge 2.0 [60]. TGV-QSM has an open-source implementation in Python [28], making it conveniently accessible for high-performance computing compared to other QSM tools that use proprietary software such as MATLAB.

QSM reconstruction is achieved using a *two-pass QSM* procedure that performs two parallel QSM reconstructions per echo, with the first including more reliable susceptibility sources, and the second including reliable as well as less reliable sources (see figure 2). The first reconstruction uses a mask produced by thresholding and binarising the magnitude signal. Thresholds can be chosen for each dataset by testing against a single acquisition, tweaking the threshold to minimise the appearance of artefacts in the final result (see figure 3). The thresholding operation automatically excludes areas with short *T*2* times, including strong and artefact-inducing susceptibility sources. The second reconstruction uses the same set of masks after applying morphological operations to fill any gaps from the first set, enabling less reliable sources to be included. An echo-dependent brain mask may also be incorporated to assist with the filling operation. A combined QSM image is then produced by filling gaps from the first QSM image using the second. After a combined QSM image is reconstructed for each echo, a final weighted average is produced by summing the combined QSMs and dividing them by the pixel-wise sum of the filled masks. By separating reliable from less reliable susceptibility sources, the two-pass technique aims to provide a substantial reduction in streaking artefacts. As an alternative to the signal magnitude, the *spatial phase coherence* can also be used as an input to the thresholding operation. Since a reduced signal in the magnitude image may not necessarily reflect susceptibility-induced dephasing, a mask based on the phase coherence may have a theoretical advantage for QSM. Spatial phase coherence is calculated by computing the second-order spatial phase gradients in three dimensions, and using a weighted combination as described in the phase-unwrapping method, Rapid Opensource Minimum Spanning TreE AlgOrithm (ROMEO) [61].

**Figure 2:**
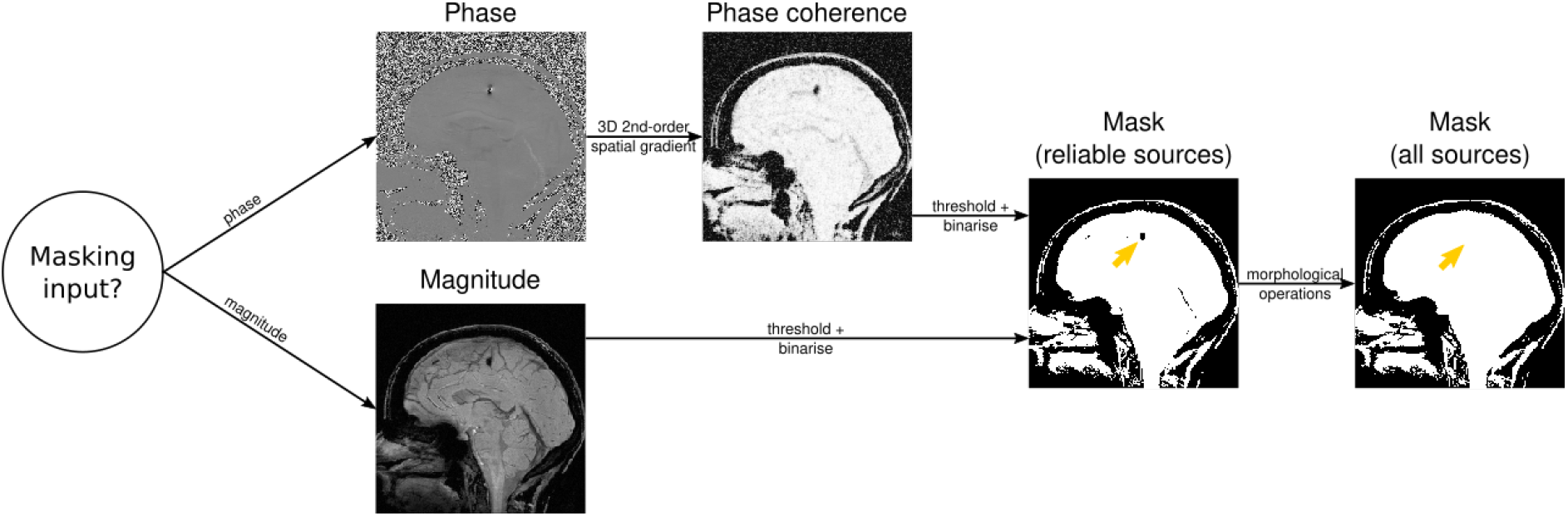
Two-pass QSM masking process for one subject with one echo. A mask is initially produced by thresholding either the magnitude or the spatial phase coherence. A second mask is then produced by filling in gaps using morphological operations. Both masks are then used in parallel QSM reconstruction steps before combination.

**Figure 3:**
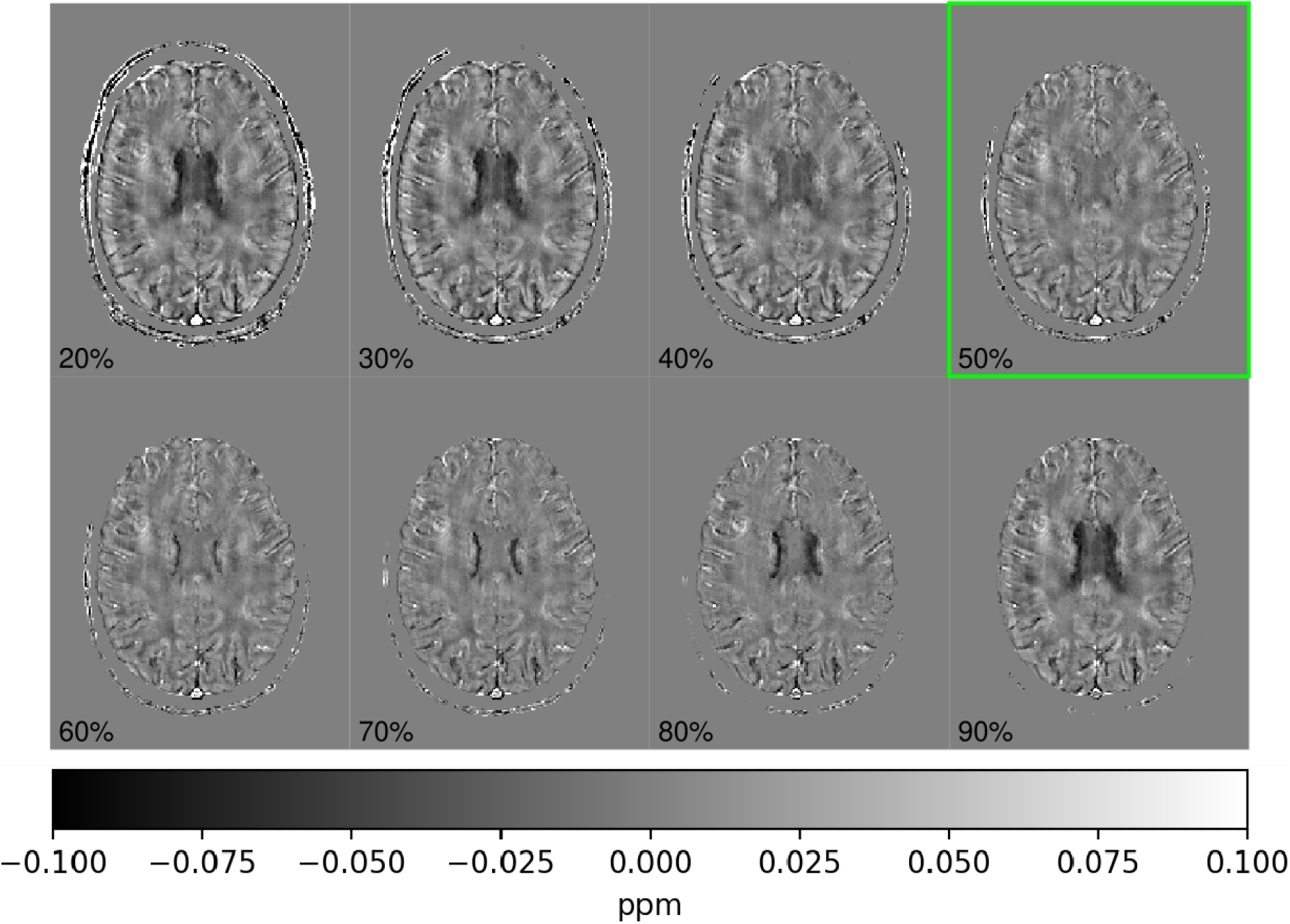
Axial QSM results from the healthy brain dataset (3 T; multi-echo) calculated using two-pass QSM with a range of magnitude signal thresholds. The slice shown is above the lateral ventricles. At some thresholds, dark artefacts are visible towards the centre of the brain caused by the dephased signal in the lateral ventricles. At a threshold of 50%, the artefacts are kept to a minimum.

Two-pass QSM was compared to the standard approach of brain masking, which applies BET [30] to the first-echo magnitude image to produce a brain mask. QSM is then reconstructed for each echo using the mask before averaging. BET was run with robust brain centre estimation enabled (to run BET iteratively) and with other settings left at their defaults. To improve performance in 7 T data, a homogeneity correction was applied to the magnitude prior to masking [56, 62].

### 2.4 Simulation

Since the combination step in two-pass QSM requires adding values into areas with low signal, a simulation was conducted to assess reconstruction accuracy in these areas. For this simulation, a dataset from the QSM Challenge 2.0 was used [41]. A spherical intra-hemispheric source was introduced to the ground truth QSM with a radius of 5 mm and a uniform susceptibility. The ground truth frequency was computed by multiplying the ground truth QSM by the Fourier space dipole kernel, to which the TGV-QSM algorithm was then applied. The experiment was performed 41 times, varying the susceptibility of the source in increments of 0.025 ppm in the range [−0.5, +0.5] ppm. The susceptibilities estimated by TGV-QSM were then measured and compared with the ground truth.

### 2.5 Template building

QSMxT generates magnitude and QSM templates using an implementation of a minimum deformation averaging method for atlas construction called Volgenmodel [58]. A template is initially constructed using the magnitude images from all subjects, and the estimated transformations are subsequently applied to the susceptibility maps to produce a QSM template. This allows for assessments of the quality of QSM results, or to compare the effects of different reconstruction techniques across large groups. QSM templates were generated for the healthy brain dataset for each masking technique to assess the group-level effects of each.

### 2.6 Segmentation and quantitative analysis

If T1-weighted images are available, QSMxT segments these using FreeSurfer via their Aseg probabilistic atlas [44–46] and registers the T1-weighted image to the magnitude in the QSM space using the MINC toolkit [63]. QSM statistics for each ROI are then exported. This segmentation and analysis pipeline was tested using the healthy brain dataset for quantitative analysis across a large group.

## 3 Results

The simulation involving lesions of various uniform strengths was conducted to assess TGV-QSM’s reconstruction accuracy with a range of simulated lesions (see figure 4). Reconstruction accuracy within the lesions was never overestimated, and the centre voxel consistently matched the ground truth. The level of susceptibility underestimation was found to correlate with the strength of the source.

**Figure 4:**
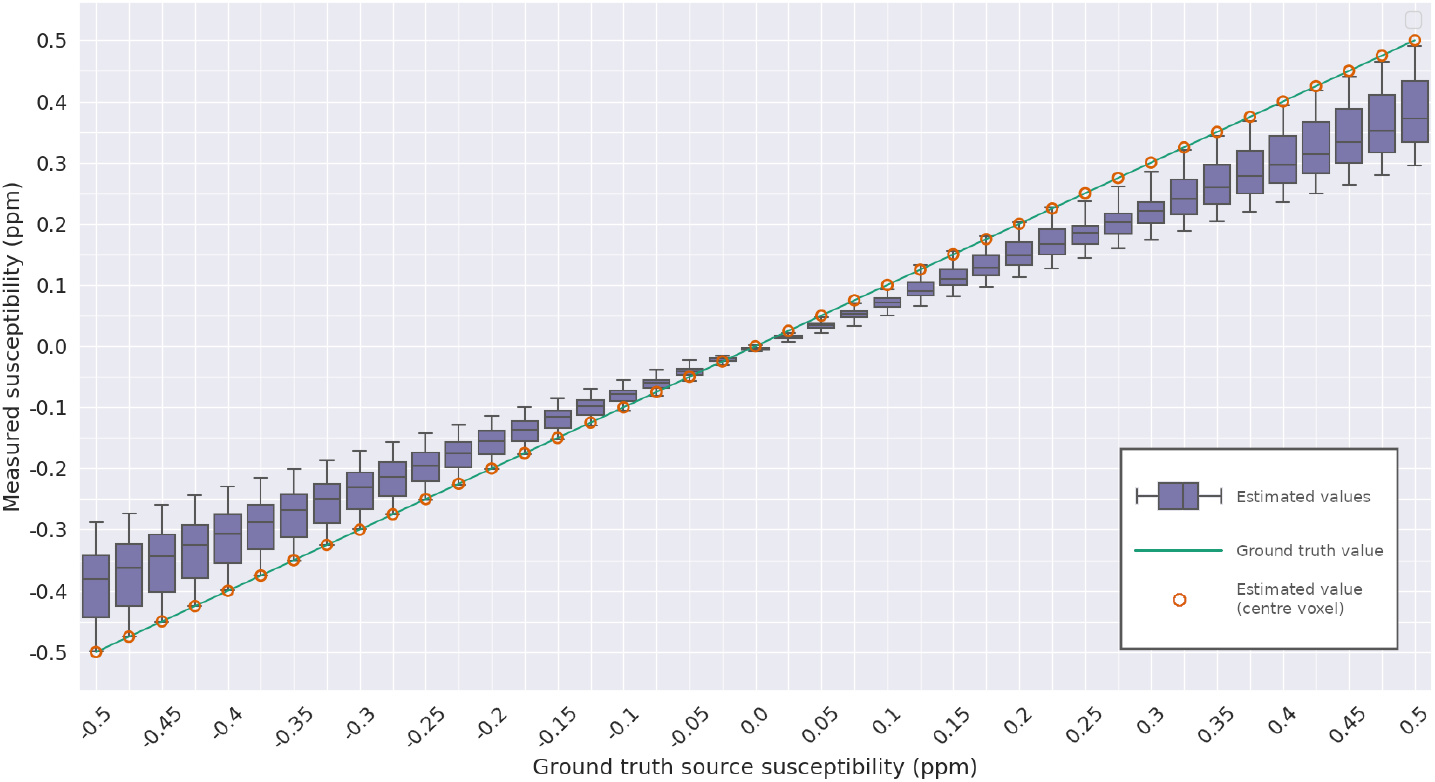
Accuracy of *TGV-QSM* in estimating the susceptibility of simulated spherical intra-cranial lesions, with their uniform strength varying between simulations. The green trendline is the ground truth susceptibility of the lesion. The orange dots are the estimated susceptibilities at the centre of the lesion. The box plots are the estimated susceptibilities across the whole lesion, extending from the lower to upper quartile values of the data, with a line at the median. The whiskers extend a maximum of 1.5× beyond the interquartile range.

Two-pass QSM was first tested in vitro using the gel phantom dataset (see figure 5). The strong susceptibility sources in the phantom caused significant streaking when all sources were included in the inversion. However, all visible streaking was successfully mitigated after the two-pass combination step. This experiment also demonstrates the masking technique’s applicability in non-brain data, as anatomical priors were not required to reconstruct QSM for the full ROI.

**Figure 5:**
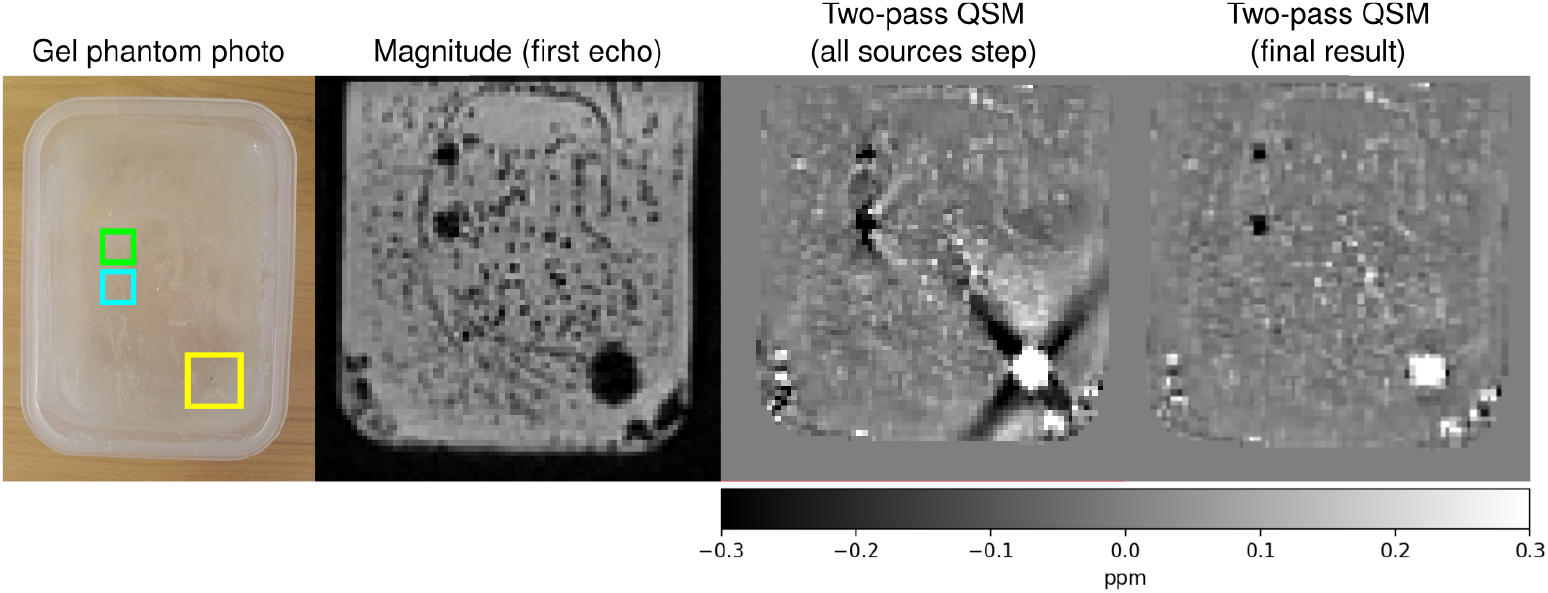
QSM results using the gel phantom dataset (3 T; multi-echo). On the left is a photo of the gel phantom containing a Gold Anchor MR+™ with 1.5% iron content (yellow square), a pure gold fiducial marker (blue square), and a tooth piece (green square). The latter three images depict the same axial slice from the magnitude, as well as intermediate and final two-pass QSM results. The final two-pass QSM image has a clear reduction of streaking artefacts surrounding the strong sources.

The QSM templates generated using the healthy brain dataset were used to examine the systematic, group-level influence of the masking technique on QSM results (see figure 6). Two-pass QSM systematically reduces the streaking that occurs near the brain boundary, particularly around the superior and transverse sagittal sinuses, with dipole-shaped reductions of artefacts visible when compared to brain masking.

**Figure 6:**
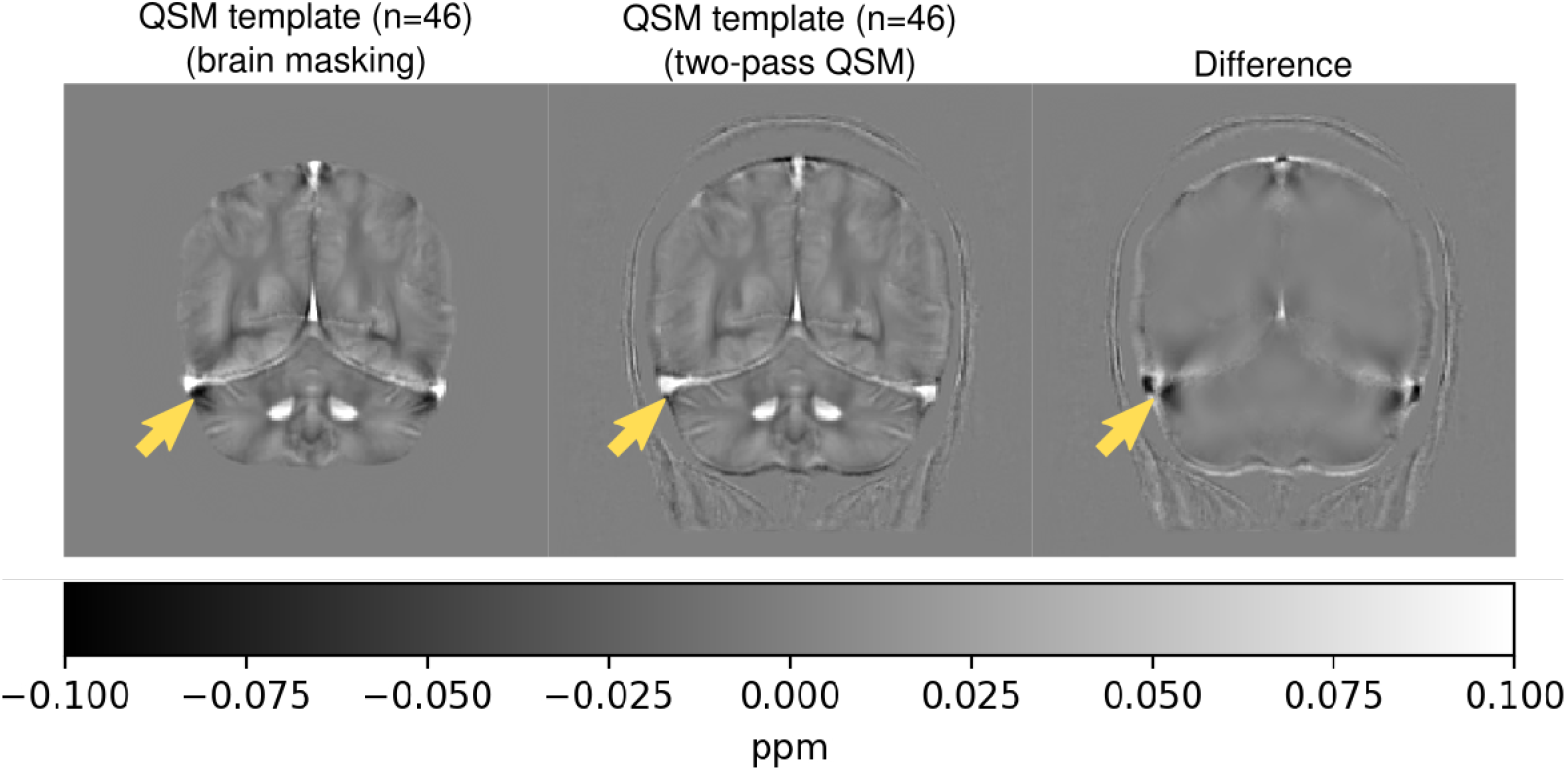
QSM templates generated using the healthy brain dataset (3 T; multi-echo; n=46); coronal slice. When brain masking is used, strong artefacts near the transverse and superior sagittal sinuses are visible (left; arrow). The artefacts are mitigated using two-pass QSM (centre; arrow), with dipole-shaped streaking visible in the difference image (right; arrow).

The QSM challenge 2.0 dataset was used to assess the effects of streaking in an individual subject and with the availability of a ground truth (see figure 7). Two-pass QSM scored significantly better on the calcification streaking metrics and overall error compared to brain masking. Further, when the threshold-based masking strategy was used, more complete anatomical detail was included in the brainstem.

**Figure 7:**
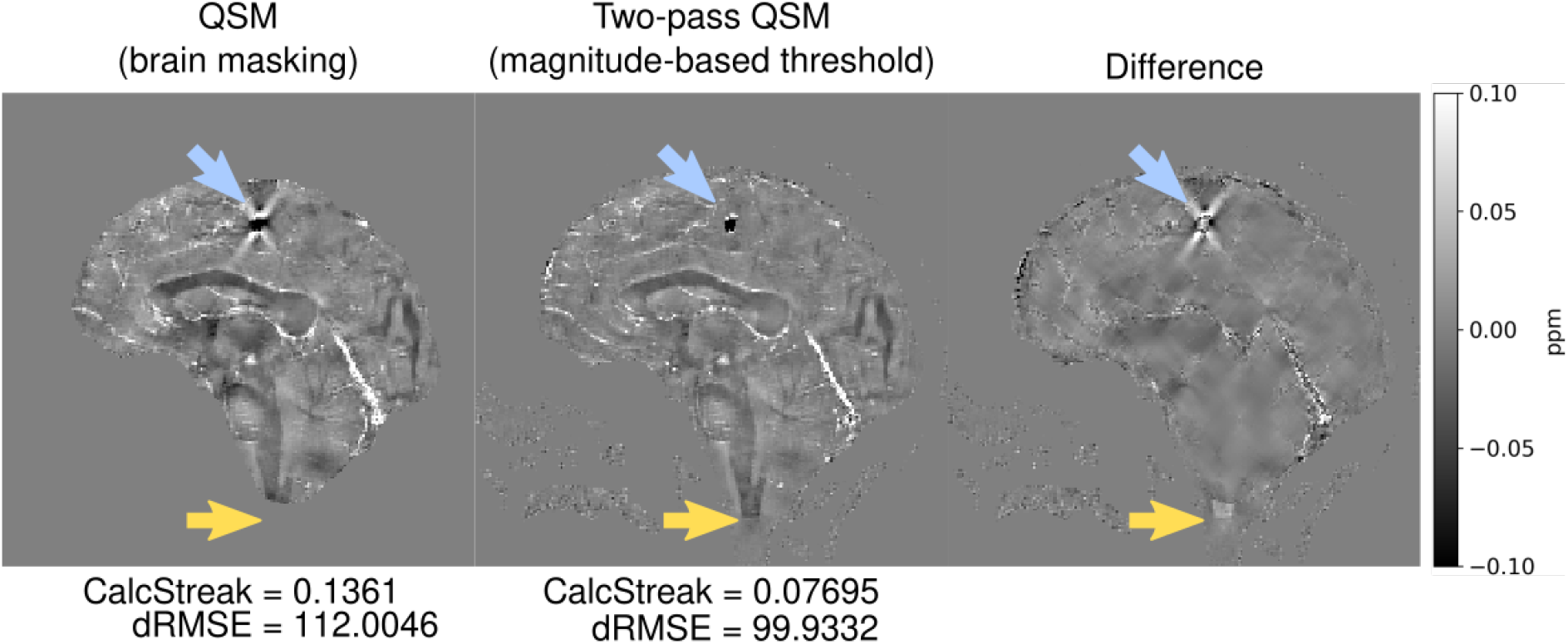
QSM results using the QSM Challenge 2.0 dataset (7 T; multi-echo; simulated); sagittal slice. Two-pass QSM reduces streaking artefacts near the brain lesion (blue arrow) and includes more anatomical detail in the brainstem (yellow arrow). The error surrounding the calcification (CalcStreak) and overall detrended RMSE across the brain (dRMSE) is notated for each method with respect to the simulated QSM challenge 2.0 ground truth (not displayed).

The brain tumour dataset was used to assess the performance of two-pass QSM at 7 T and in clinical cases. In this dataset, BET regularly failed to mask the whole brain, sometimes missing crucial tumour detail (see figure 8). Further, two-pass QSM mitigated the artefacts surrounding tumours, veins and vessels, generating more plausible QSM values (see figures 8 and 9).

**Figure 8:**
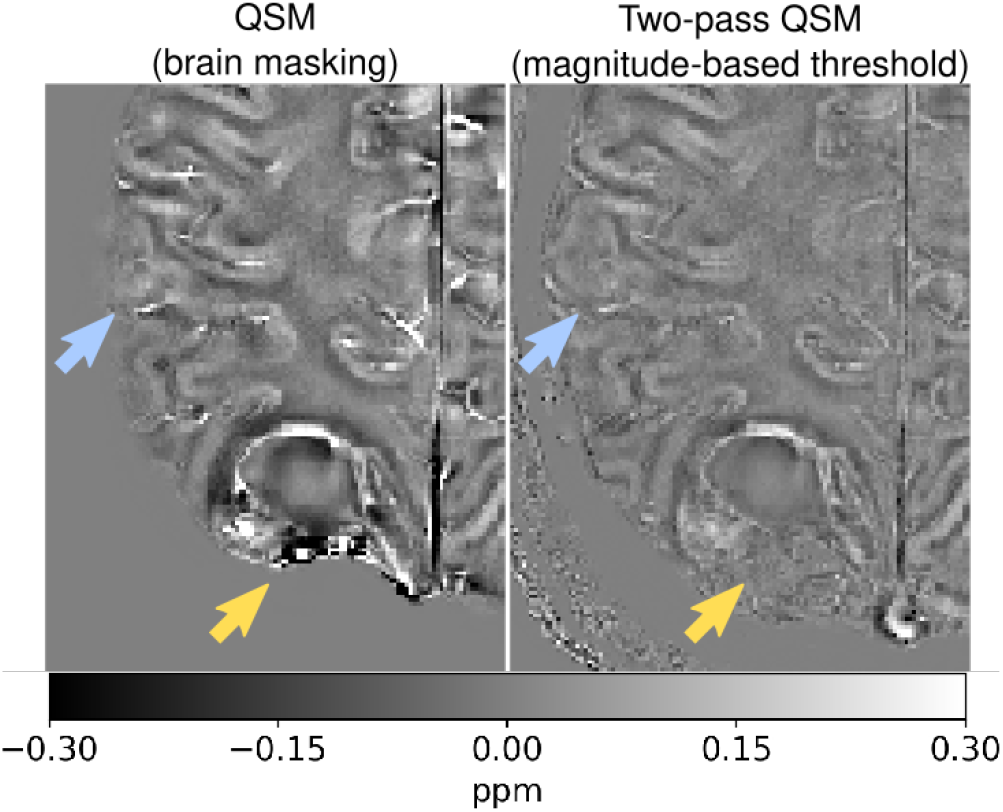
QSM results from the brain tumour dataset (7 T; multi-echo) computed using brain masking and two-pass QSM; axial slice region. The brain mask excluded part of the tumour included by QSMxT (yellow arrows). Our proposed technique also included more cortical detail (blue arrows).

**Figure 9:**
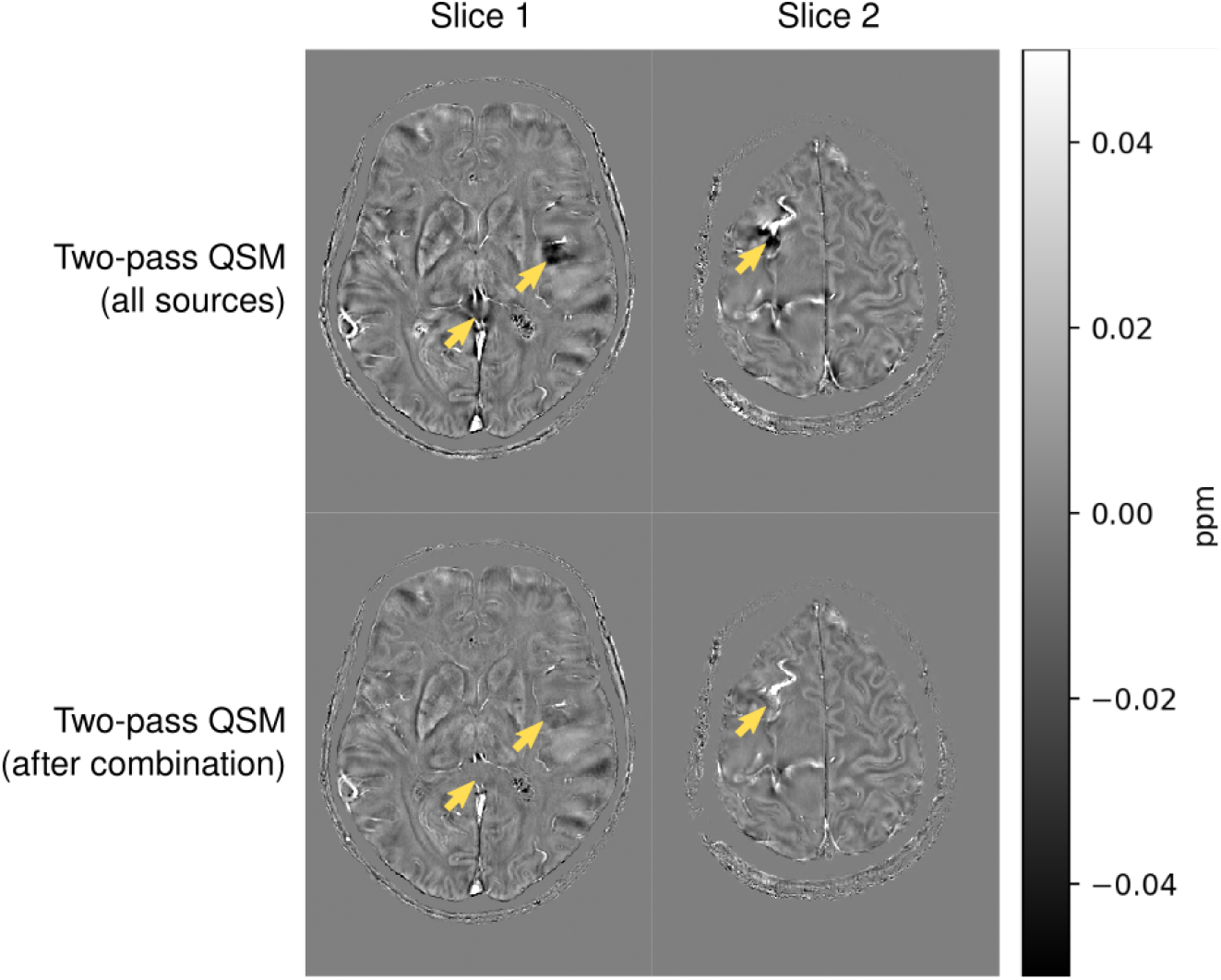
QSM results from the brain tumour dataset (7 T; multi-echo) computed using two-pass QSM, before and after the combination step; two axial slices. Significant reductions of artefacts can be seen near vessels after two-pass combination (yellow arrows).

## 4 Discussion and conclusions

A novel two-pass masking and QSM reconstruction strategy has been presented that separates susceptibility sources using masks based on the MRI signal. Compared with standard brain masking, the threshold-based technique includes more anatomical detail without priors, especially in pathology (see figures 7 and 8), facilitating studies in tumour patients and more widely across the head and neck. The two-pass reconstruction mitigates the streaking artefacts caused by high dynamic ranges of susceptibility values and signal decay with TE (see figures 5, 6, 7, 8, and 9). The improved accuracy in the QSM challenge 2.0 metrics confirm that the artefact reductions lead to a more accurate QSM reconstruction (see figure 7). The reduction of artefacts also allows the boundaries of strong sources, including vessels and veins, to be depicted with greater clarity, facilitating object identification, segmentation and quantitative analysis. Further, the two-pass QSM technique and its masking strategies are automated within the open-source QSM framework, QSMxT, which was used for all results presented with default parameters across all datasets. This demonstrates the robustness of the technique across institutions, field strengths, scanners and acquisitions. Finally, the conceptual simplicity of the thresholding operation makes it straightforward to implement in conjunction with any QSM algorithm requiring a mask, and it can also run in a single pass if computational and time resources are limited.

A limitation of the two-pass results presented is the lack of a ground truth, which was only available for the QSM challenge dataset and for the simulated lesions. Therefore, the claim that the artefact reductions lead to a more accurate QSM generally is based on the QSM challenge dataset, which uses a single field strength and imaging resolution. Further, the robustness of the two-pass technique and invariance to anatomy is based on results across multiple imaging resolutions, field strengths, pathology, and in simulations and phantoms. Robustness in QSM applications more broadly across the human body or in other animals is yet to be investigated.

Several other multi-stage QSM reconstruction techniques exist with similar artefact mitigation results, including QSMART [35], STAR-QSM [34], and SDI [36]. However, the two-pass QSM technique provides two novel features: first, it does not require anatomical priors for brain masking or source separation, including optimised and application-specific filters, making it potentially useful across a much wider array of applications; secondly, it requires minimal to no parameter tuning across applications and datasets, making automation over such datasets practical, with the potential to be used in clinical environments.

As two-pass QSM involves inserting values with low SNR, it was necessary to assess the reliability of *TGV-QSM* in these areas with respect to a ground truth (see figure 4). The simulation results indicate that the spread and accuracy of the reconstructed values negatively correlates with increasing source intensity. This systematic underestimation of susceptibility values has been previously reported to be caused by imaging resolution and signal strength [64]. The single strongest value at the centre of the source was found to be a very accurate representation of the source overall. Critically, the simulations demonstrate that all simulated sources can be distinguished from one another based on their estimated susceptibilities, indicating that the values inserted using two-pass QSM may be analysed and compared. It should be noted that the introduced source in the simulation had a uniform intensity and spherical shape, and the simulation did not include a background field, signal noise, phase wraps, or temporal evolution.

To make the two-pass technique accessible and reproducible in the QSM community, it was integrated into the new QSM framework, QSMxT. Existing QSM processing frameworks have limited flexibility and scalability, with restrictive algorithm selection or programming language requirements, and with most tools only able to process one subject at a time, and with manual intervention required between steps. Further, the lack of QSM tools with integrated segmentation and analysis hampers the use of QSM among researchers, especially those less experienced with segmentation and quantitative MRI. QSMxT requires only a single dependency to run, is compatible with every operating system, and has no restrictions on algorithm selection, with the full NiPype pipeline easily configurable. QSMxT can also take advantage of HPC systems to process many datasets and computational steps in parallel. With a single command, QSMxT can run one of several pipelines to standardise input data, reconstruct multi-echo susceptibility maps using two-pass QSM, generate a group space along with QSM and magnitude templates, and provide full segmentations and quantitative outputs. QSMxT significantly lowers QSM’s barrier to entry, providing researchers with a complete set of tools to include QSM in their studies.

## 5 Data Availability Statement

We facilitate the reproducibility of our study by providing the full implementation of the pipeline (https://github.com/QSMxT/QSMxT, commit hash 6308062). The pipeline can be executed in NeuroDesk (https://github.com/NeuroDesk).

## 7 Ethics approval

The study data included was approved by the relevant ethics committees; the 7 T tumour study data by the ethics committee of the Medical University of Vienna, and the 3 T healthy brain study data by the ethics committee of the South Australian Health and Medical Research Institute (SAHMRI).

## 8 Acknowledgements

We thank the participants involved in this study. MB acknowledges funding from Australian Research Council Future Fellowship grant FT140100865 and SR from the Marie Skłodowska-Curie Action MS-fMRI-QSM 794298. This research was funded by the Australian Research Council Training Centre for Innovation in Biomedical Imaging Technology (project number IC170100035) and funded by the Australian Government. Additional support was provided by the Austrian Science Fund (FWF): 31452 and KLI-646. The authors acknowledge the facilities and scientific and technical assistance of the National Imaging Facility, a National Collaborative Research Infrastructure Strategy (NCRIS) capability, at the Centre for Advanced Imaging, the University of Queensland. We wish to acknowledge QCIF for its support in this research by providing high performance computing and storage resources.

